# DirtyGenes: significance testing for gene or bacterial population composition changes using the Dirichlet distribution

**DOI:** 10.1101/204321

**Authors:** Laurence M Shaw, Adam Blanchard, Qinglin Chen, Xinli An, Sabine Tötemeyer, Yong-Guan Zhu, Dov J Stekel

## Abstract

High throughput sequencing, and quantitative polymerase chain reaction (qPCR), can detect changes in bacterial communities or the genes that they carry, between different environments or treatments. These methods are applied widely to microbiomes in humans, animals, soil and water; an important application is for changes in antimicrobial resistance genes (ARGs). However, at present, there is no statistical method to determine whether observed changes in the overall composition are significant, or result from random variations between samples. Therefore researchers are limited to graphical descriptions. We describe a novel statistical method to determine whether or not observed differences in bacterial populations or their genes are significant. It can be used with data from shotgun metagenomics, 16S characterisation or qPCR. It can also be used for experimental design, to calculate the number of samples needed in future experiments. We show its application to two example data sets. The first is published data on bacterial communities and ARGs in the environment, in which we show that there are significant changes in both ARG and community composition. The second is a new data set on seasonality in bacterial communities and ARGs in hooves from four sheep. While the observed differences are not significant, we show that a minimum group size of eight sheep in a future experiment would provide sufficient power to observe significant changes, should the already observed changes be true. This method has broad uses for statistical testing and experimental design in experiments on changing microbiomes, including for studies on antimicrobial resistance.

## INTRODUCTION

Bacteria live in complex communities, whether in water, in soil, or on larger organisms, as the microbiota of organs such as the gut or the skin. New high throughout technologies, including high throughout sequencing (Venter et al. 2004), or qPCR arrays (Looft et al. 2012), allow for the characterisation of microbial communities (Truong et al. 2015), or the genes that they carry (Zhu et al. 2013). This has broad application across biomedical and environmental science, and in particular, allows for detection of changes in communities or the genes that they carry in the face of biological (Marti et al. 2017), chemical (Chambers et al. 2015) or environmental (Garner et al. 2016) factors. Of particular relevance are studies on antimicrobial resistance (Zhu et al. 2013; Su et al. 2015; Chambers et al. 2015; Xiao et al. 2016; Chen et al. 2016; Garner et al. 2016), as well as studies in other areas, such as changes in gut flora in different human communities (Shankar et al. 2017) or through the ingestion of probiotics (Unno et al. 2015).

For robust reporting of research, it is important to use statistics correctly to ascertain whether there is evidence that the observed changes in the taxonomic or genetic composition of a community reflect the factors under study, as opposed to merely reflecting random variation between the samples. By *composition*, we refer to the proportions of individual taxanomic or gene classes within the population; this can be contrasted with *abundance*, which refers to the number of individuals, either overall, or of specific taxanomic or gene classes, or *overall structure*, which takes into account both abundance and composition. However, much previous work, especially in antimicrobial resistance, has largely focussed on visual methods for analysing data, using tables and graphical representations to compare overall population compositions or structures (Zhu et al. 2013; Wang et al. 2014; Chambers et al. 2015; Xiao et al. 2016; Garner et al. 2016), without providing statistical support to evidence change. Methods such as taking diversity indices or principal coordinate analysis (Chen et al. 2016) have allowed for a more in depth analysis of population structures than using pie charts/compositional bar charts. However, these techniques are also visual, and do not have a clear blueprint to determine whether observed differences in structure are significant.

A related problem is experimental design. Power analyses are routinely used to determine how many individuals to recruit into experimental studies. However, power analyses require a given statistical method for analysis of the resulting data. In the absence of a statistical test for difference in taxanomic or genetic composition, there is no rational way to determine how many individuals to recruit into a study.

There are methods available to answer similar questions. For example, methods developed for cDNA library (Stekel et al. 2000) or RNA-seq analysis (Robinson et al. 2010; Anders and Huber 2010; Hardcastle and Kelly 2010), though not commonly used in this context, could be applied to metagenomics data on a taxon-by-taxon or gene-class-by-gene-class basis to identify individual taxa or gene classes that are significantly different. While such analysis could be of value, it does not answer the question about whether the overall community has changed. Moreover, such analyses would be subject to considerable numbers of false positives because of the number of tests applied, and would be difficult to correct for multiple testing because of the high level of correlation between different taxanomic or gene classes.

In this work, we develop a statistical test to determine whether the taxa-nomic or gene composition of microbial community samples are statistically different between two or more sets of treatments or conditions. By considering the composition of ARG or bacterial population data in terms of a *Dirichlet distrubution* (Maier 2014), we are able to perform a likelihood ratio test in order to obtain a p-value to determine the level of evidence that population compositions vary significantly across multiple environments. We also outline a goodness-of-fit test to determine whether the Dirichlet distribution provides a reasonable model for a given dataset, without which the significance test outlined would not be valid. Moreover, we show how this test can be used for experimental design, in order to identify numbers of individuals to use in an experiment. As examples, we apply the method to two data sets: soil microbiome data following manure amendment (Chen et al. 2016), that uses qPCR arrays and 16S metagenomics; and previously unpublished data from a pilot study on seasonal changes in the microbiota of ovine hooves, that uses shotgun metagenomics. The interdigital skin environment of cloven hoofed animals has a dynamic bacterial community, composed mainly of skin and faecal colonising bacteria, but also soil microbes (Maboni et al. 2017; Zinicola et al. 2015). These exemplify the broad applicability of this approach.

## RESULTS

We first briefly describe the test statistic used to compare populations across multiple environments, before conducting two example analyses. A more detailed derivation of the test statistic is given in the Methods section which can be applied to any multitype population, including bacterial populations.

Consider the profiles of samples taken from a set of different treatments or environments with replication. These profiles could be proportions of taxanomic groups (at any level) or proportion of gene groups, for example classes of antibiotic resistance genes. We test the null hypothesis, which states that any differences in composition between treatments / environments are a result of pure chance, against the alternative hypothesis that the population composition is affected by the treatment / environment.

We classify the composition of the taxanomic or genetic groups by a *Dirichlet distribution*. Under the null hypothesis, the composition of every sample is drawn from the same Dirichlet distribution. Under the alternative hypothesis, the parameterisation of the Dirichlet distribution governing the composition of the groups changes according to the treatment or environment.

The test statistic takes the form

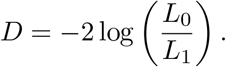

where *L*_0_ and *L*_1_ are the maximum values of the likelihood function for the Dirichlet distribution parameterisation under the null and alternative hypotheses respectively. In the limit of a large number of samples, under the null hypothesis, *D* would follow a 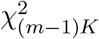 distribution where *m* is the number of environments in the study and *K* is the number of ARG classes. However, most experiments do not have sufficient replication for this approximation to be valid. Therefore we use a randomization procedure to determine p-values. Further details on the derivation and calculation of the test statistic, and of the randomization procedure, are in the Methods section.

### Example Analysis: Sewage Sludge, China

Our first data set considers the effect of the application of sewage sludge to soil (Chen et al. 2016). ARG and bacteria populations were profiled under eight different soil treatments with three samples being taken from each soil type (giving a total of 24 samples). Following the labelling of Chen *et al.* 2016, the soil types are referred to as follows: *CK* is a control soil containing no manure or urea (a chemical fertiliser); *0.5N* and *1N* soils contained urea with double the application rate in the 1N soil; *CM* contained chicken manure; while *0.5SS, 1SS, 2SS* and *4SS* contained sewage sludge with the application rate doubling with the coefficient. All the manure soils also contained the same application of urea as 0.5N.

Figure 1 displays the compositional data of the ARGs by the class of drug to which they are resistant and bacteria by phyla for each of the samples from these data. Note that results from one of the 4SS samples was missing from the bacterial data. In each case, only ARG classes/bacterial phyla that were ever-present across all samples and accounted for at least 1% of the population in at least one sample were included. Classes/phyla not meeting these criteria were aggregated into a group called *LRT other*. The reasons for this are outlined in the Methods section.

Visual inspection of the plots in Figure 1 suggests that the composition of ARGs by class shows little variation within the same soil types but does seem to change with the environment. However, it is not clear from the figure whether the soil environment has an affect on the composition of bacterial populations.

Applying our likelihood ratio test to the data gave p-values of 1.03 × 10^−125^ for the ARG data and 1.32 × 10^−19^ for the bacterial data using the χ^2^ method. These results were validated using the randomization approach (5000 random samples) which found p-values of < 0.0002 and 0.0266 for the ARG and bacterial data respectively.

This provides strong evidence to suggest that the composition of both ARG and bacteria populations are affected by soil amendment; the evidence is particularly strong for the ARG data. Goodness-of-fit testing supports the notion that the Dirichlet distribution provides an adequate model for both sets of compositional data, with p-values of 0.55 for the ARG data 0.479 for the bacteria data, using 10000 simulations. Note that it is these non-significant p-values that indicate that the fit is good. Details on how goodness-of-fit testing is performed for these data can be found in the Methods section.

**Figure 1:**
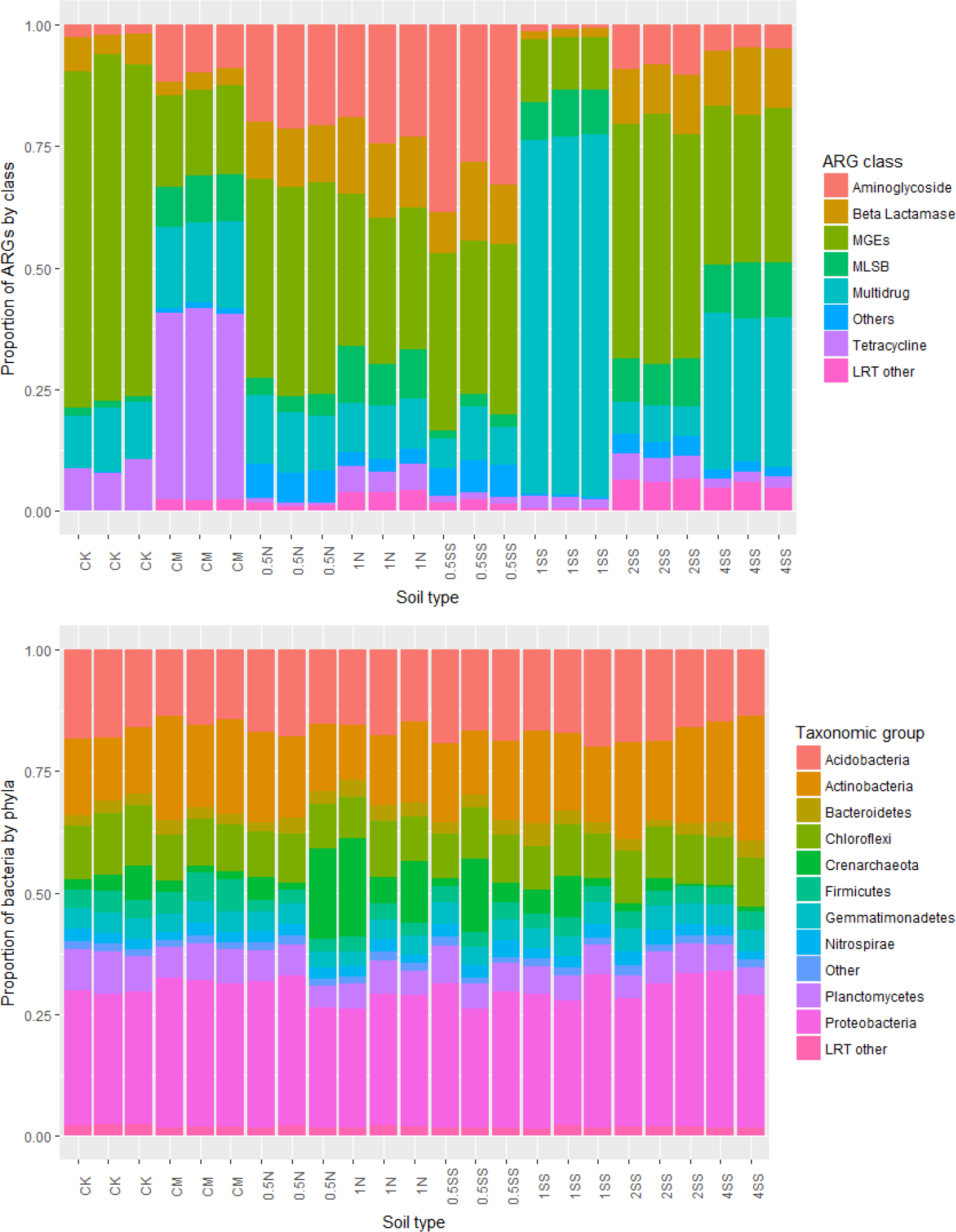
Compositional data of ARGs by class and bacterial taxa by phyla for each of the soil samples

### Sheep Hooves, UK

The second dataset looks at how sheep hooves provide a reservoir for the spread of AMR. Four different sheep were used and the ARG and bacterial populations on their hooves were profiled in three different seasons (Winter, Spring and Summer), giving 12 samples overall. Further details on how the data were collected are given in the Methods section.

The compositional data of both the ARG and bacterial populations for each sample is shown in Figure 2. Again, all classes/phyla that were not ever-present and did not account for at least 1% of at least one sample were amalgamated into the *LRT other* category.

For these data, visual inspection suggests that changing seasons does not explain much of the variation in either the ARG or bacterial population data. The Dirichlet likelihood ratio test gave p-values of 0.057 for the ARG data and 0.597 for the bacteria data using the chi-squared method. Randomization testing gave p-values of 0.586 and 0.898 respectively. In both cases the null hypothesis that season does not have a significant effect on population composition would not be rejected at the 5% level. Goodness-of-fit testing gave p-values of 0.484 and 0.511 for the ARG and bacterial data respectively, again suggesting that our proposed Dirichlet model provides an adequate fit to these data.

**Figure 2:**
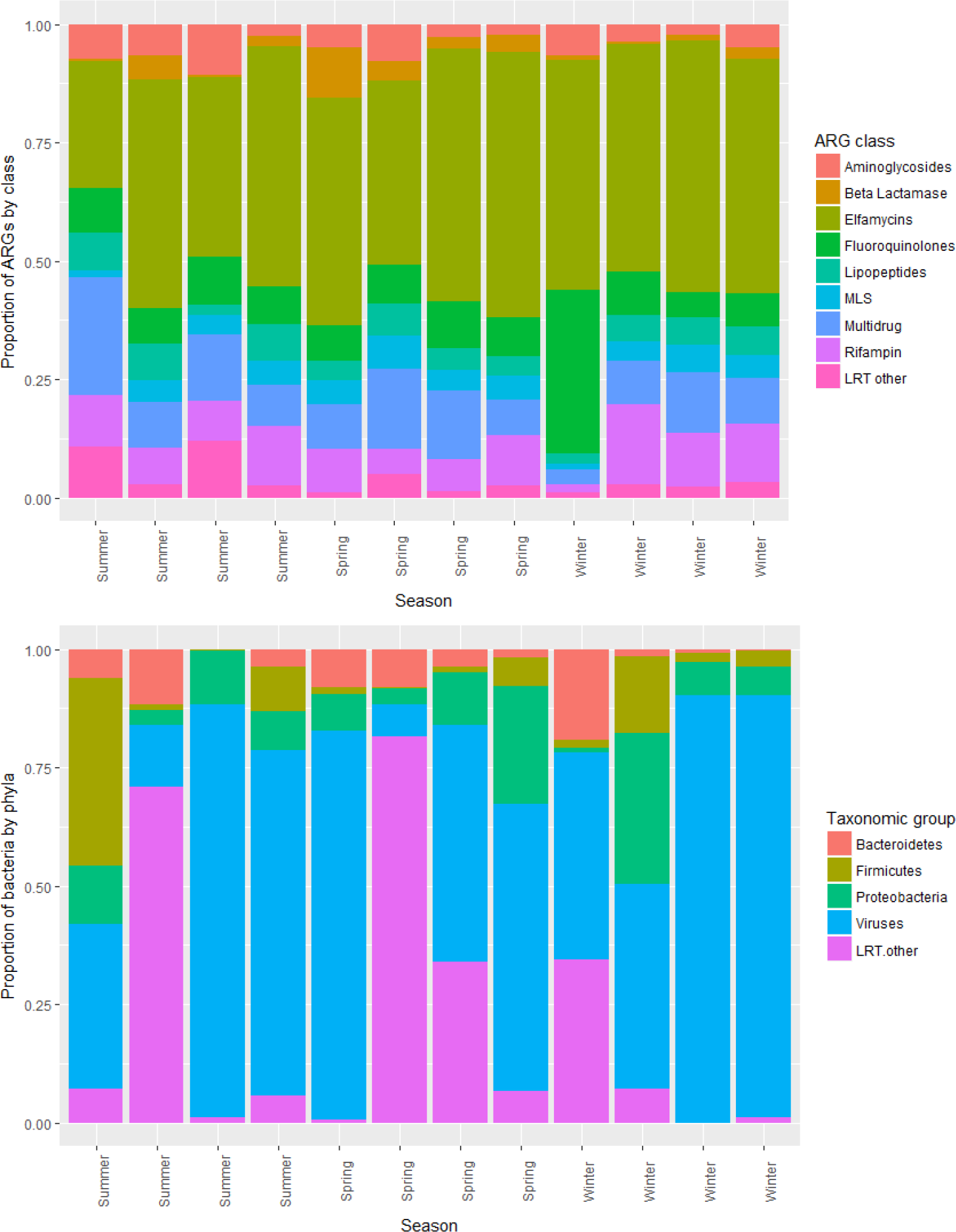
Compositional data of ARGs by class and bacterial taxa by phyla for each of the sheep hoof samples

### Power Analysis

The test method can also be used for experimental design, through its use in power analysis to estimate group sizes in future experiments. This is achieved by assuming the alternative hypothesis that population composition is affected by the treatment or environment is true, and using the currently observed data as the *best guess* for the distribution of the population in each environment. Full details are given in the methods section.

In the sheep hoof pilot study, we could not find significant evidence to support the hypothesis that ARG or bacterial population structures are affected by season; however, the pilot study from which the data derive only used a group size of four sheep for each season. Assuming that there are differences between population structures in the three seasons, Figure 3 shows estimates the power of future tests for groups containing up to 8 sheep for both the ARG and bacteria data. The power estimates were generated using 500 simulations for each group size.

We observe that if there are differences between the ARG populations then future experiments should be very powerful at the 5% level regardless of the number of sheep used. (Recall the p-value close to 0.05 for the original data under the χ^2^ test.) Creating an experiment with sufficient power to observe differences in bacterial populations between seasons requires more sheep: an experiment with a group of eight sheep should give us more than an 80% chance of finding significant evidence to support such a claim.

**Figure 3:**
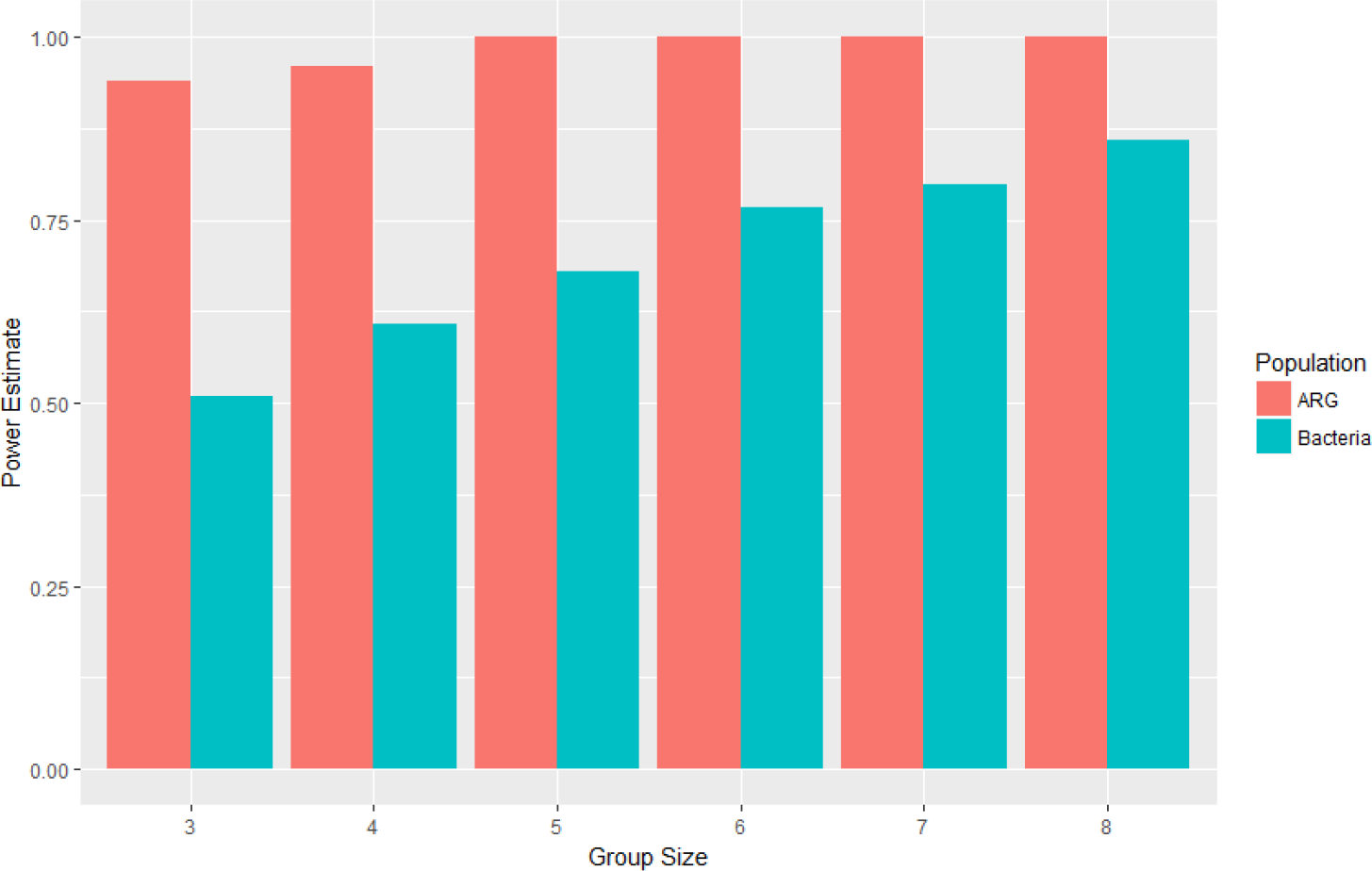
Power estimates of the Dirichlet LRT test for repeated experiments of the sheep hooves experiment with different numbers of sheep

## DISCUSSION

We have described and demonstrated a method for comparing gene or tax-anomic composition of populations across a range of treatments or environments by modelling compositional population data in terms of a Dirichlet distribution and using a likelihood ratio test. Methods for assessing whether the selected model provides a reasonable fit to the data, without which the test would not be valid, and also a procedure for estimating the power of future experiments, have also been outlined. These provide new and valuable analyses that can be used both for assessing evidence from microbial communities studies, and for estimating how many samples to use in future studies.

Our primary motivation and application has been to antimicrobial resistance, although the method is valid for any multi-type microbial community. We have built on previous contributions to this field by describing differences in entire populations using a single test statistic, rather than relyng on analysing one ARG/bacterial taxa at a time between different environments. This allows us to draw quick inferences as to whether changing environments are having an effect on the composition of our data (by simply reading off and interpreting a p-value) and removes the need to consider problems associated with making multiple comparisons (Stekel et al. 2000).

An important limitation of the test is to only include population types or classes that are present across all samples in the data, and that meet the criterion that the type accounts for at least 1% of the population in at least one of the samples. All types not meeting both criterion need to be aggregated into another class. This is essential for the method to give reliable estimates for the parameters of the Dirichlet distributions. In most cases, the interest is in the differences between the more abundant classes within the data, and so this limitation is unlikely to have a major impact on interpretationo of the data. The issues behind the limitation are explored in greater detail in the Further Considerations part of the Methods section.

The method outlined in this paper has the advantage being able to look at entire multitype populations (both in AMR and other fields) but does not use information on abundances, which may also provide an insight into differences between environments. A possible extension to this method would be to model the overall structure of the observed populations using either a multinomial or Dirichlet-multinomial distribution. Such a distribution would be associated with a probability vector ***p*** (which may be fixed or be drawn from a Dirichlet distribution for each observation) denoting the probabilities of a random individual drawn from the population belonging to each type. The distribution would also need a number of trials, *N*, representing to overall size of the population, which would also be a random variable based on the distribution of the overall population sizes.

One would need to decide upon an appropriate distribution for *N* and also consider the issue that populations with larger *N* would contribute more to the likelihood function, meaning that each observation would not make an equal contribution to the likelihood ratio test statistic. However, such a method would use all of the information available from a multitype population and would remove the problem needing ever-present types since zeros are supported within multinomial distributions. Instead, classes would only need to be present in at least one sample from each environment.

For potential users of the test, we have produced R code, both as a Supplementary File, and on the FigShare data sharing web site, together with details of how to use the functions. Moreover, we foresee considerable potential for inclusion of this method into pipelines for analysis of antimicrobial resistance data (Arango-Argoty et al. 2016; Yang et al. 2016).

## METHODS

### Derivation and considerations of the test statistic

We describe how to model multitype population compositions in terms of a Dirichlet distribution before explaining how the test statistic used to compare compositions is obtained. We also address some important considerations when carrying out the likelihood ratio test, including outlining our method for goodness-of-fit testing, designed to ensure that the Dirichlet distribution provides an appropriate model for the observed community compositions.

### Multitype population structures as Dirichlet distributed random variables

The Dirichlet distribution is a standard distribution in statistics in *K* > 2 dimensions which is described by the parameters ***α*** = (*α*_1_,*α*_2_,…,*α*_*K*_) with *α*_*k*_ > 0 (*k* = 1,2,…, *K*). A single, random draw from the Dirichlet distribution will give an observation of the form (*x*_1_, *x*_2_,…, *x*_*K*_) where

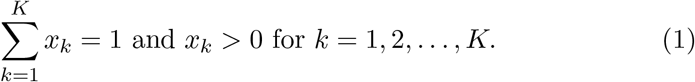

The expected value of *X*_*k*_, the *k*^*th*^ component of a random variable distributed as described above, is given by

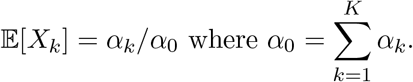

Note that the expected values do not change if each of the *α*_*k*_ parametrising the Dirichlet distribution are multiplied by some constant *c* > 0. However, the variance is affected with larger *α*_0_ values resulting in smaller variances. Specifically,

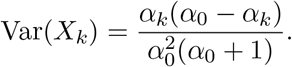

Now suppose that we have an observation of a multitype population containing *K* classes of individuals with *y*_*i*_ individuals in each class (*k* = 1,2,…, *K*). For *k* = 1,2,…, *K*, let *x*_*k*_ denote the proportion of individuals of class *k* within the population, given by

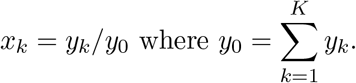

Since the *x*_*k*_ meet the conditions set out in (1), we propose that the proportions of individuals in each class can be considered to be a random draw from a Dirichlet distribution with unknown parametrisation ***α***.

### Likelihood Ratio Test

Suppose our multitype population has *K* classes and we have *m* environments of interest. For *i* = 1, 2,…, *m*, let *n*_*i*_ be the number of observations of population compositions in the *i*^*th*^ environment. Let *x*_*i*,*j*,*k*_ denote the proportion of individuals in class *k* in observation *j* from the *i*^*th*^ environment (*i* = 1,2,…, *m*; *j* = 1,2,…, *m*; *k* = 1,2,… *K*). We assume that the composition of a randomly selected population in environment *i* follows a Dirichlet distribution with parameters ***α***_*i*_ = (*α*_*i*,1_, *α*_*i*,2_,…, *α*_1,*k*_). Let α be the concatenation of the vectors ***α***_1_, ***α***_2_,…, ***α***_*m*_ and ***x*** denote the set of all observations *x*_*i,j,k*_. The likelihood of a parametrisation *α* given a set of observed population compositions *x* is given by

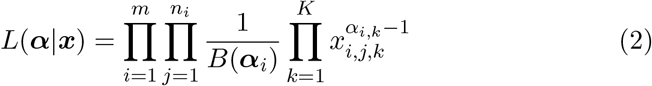

where *B*(***v***) denotes the multivariate beta function, applied to the vector *v*. We wish to test the following nested hypotheses:

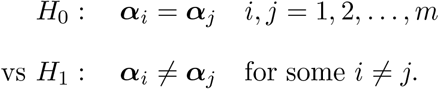

Let *L*_0_ be the restricted maximum value of *L*(***α***|***x***) under *H_0_* and *L_1_* be the unrestricted maximum value of *L*(***α***|***x***) under *H*_1_. Our test statistic is given by

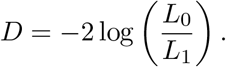

By standard statistical theory, *H*_0_, *D* follows a 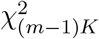 distribution, since (*m* − 1)*K* is the number of additional parameters under our alternative hypothesis.

Should a randomization procedure be required due to limited replications, as described in the Results section, this can be achieved by calculating *D* as described above for the original data. A further set of test statistics for randomized data can be calculated by randomly permuting the environments associated with the results from each of the samples in the data. The proportion of test statistics from the randomized data that are greater than the value *D* from the original data provide a p-value for the hypothesis test outlined above. Results quoted in this paper are from randomization procedures with 5000 trials.

Code for calculating the test statistic and its associated p-value in R Statistical Software (either by comparison to the relevant χ^2^ distribution or by randomization) is also given as supplementary material. Further detail on the derivation and computation of maximum likelihood estimators form a Dirichlet distribution may be found in the note by Minka (Minka 2000).

### Goodness-of-fit Testing

The likelihood ratio test outlined above determines which of two models based on the Dirichlet distribution most best describes our data. However, our test is based on the assumption that population composition can be described the Dirichlet distribution. We describe a test to determine whether the selected Dirichlet distribution model provides an adequate fit to the given data. Code for performing this test is provided as part of the likelihood ratio test code given in the supplementary material.

Standard goodness-of-fit tests such as Pearson’s chi-squared test or the Kolmogorov-Smirnov test are only applicable to one-dimensional distributions. Since the Dirichlet distributions that we are testing are *K*-dimensional (*K* > 2), such tests are inappropriate and thus we use an ad-hoc test best on likelihoods to generate a p-value for goodness-of-fit. The test compares the likelihood given the observed data to likelihoods given simulated data from the proposed distribution with the same parametrisation.

The test is designed as follows. Select a Dirichlet distribution model with parametrisation α to model the observed data. Calculate the (log) likelihood of the parametrisation given the observed data using (2) and denote this value by *T*_0_. Choose a suitably large integer *N* and simulate *N* copies of the data, using the same *K*, *m* and *n*_*i*_ (*i* = 1,2,… *m*) as in the observed dataset. For *j* = 1,2,…, *N* let *T*_*j*_ be the likelihood of ***α*** given the *j*^*th*^. simulated data set.

For this test, our null hypothesis is that our observed data come from Dirichlet distribution model with parametrisation ***α***. Under this hypothesis, the likelihood of ***α*** given the observed data, *T*_0_, should not be significantly different to the *T*_*j*_ (*j* = 1,2,…,*N*), the likelihoods of ***α*** given the simulated data. Specifically, the proportion of the *T*_*j*_ < *T*_0_ can be used as a p-value for testing the null hypothesis, since *T*_0_ will be smaller than most of the *T*_*j*_ if the proposed Dirichlet distribution provides a poor fit to the observed data.

### Power Testing

To perform a test estimating the power of future experiments we assume that the alternative hypothesis *H*_1_ is true and specifically, assume that the population compositions are described by ***α*̃**, where ***α̃*** is the value of ***α*** which maximises *L*(***α***|***x***) under *H*_1_ for our original data.

The power of a future experiment may be achieved by selecting a significance value for future tests (usually 5%), a number of replications of the experiment in each environment *n*, and, as with the goodness-of-fit testing, a suitably large integer *N* which denotes the number of simulations to run. Simulate data with *n* replications in each of the *K* environments by simulating *n* copies of each of the *K* Dirichlet distributions described by the parameterisation ***α̃***. This creates a single simulation of new data with *n* replications in each environment which can then be put through the likelihood ration test. If the resulting p-value from this test is less than the significance value selected then this is recorded as a success. The proportion of successes in *N* replications of this procedure provides an estimate of the power of a future experiment with *n* replications in each environment.

*Note that the test described here is computationally expensive. The attached R code will run much slower if the user includes power testing options*.

### Further Considerations

In our example analysis we make two important decisions which affect both the test statistic and the χ^2^ distribution that it is compared to. These are to only consider classes within the multitype population that account for at least 1% of the population in at least one of the samples and to only include classes which are present across all samples.

We are generally interested in differences between the more abundant classes in a given multitype dataset. From the derivation of the likelihood ratio test given earlier in this section, *D*, follows a 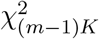 distribution under *H_0_*. Suppose *K*′ < *K* of the classes fail to meet a minimum percentage threshold such as the 1% given above. The number of degrees-of-freedom associated with the test statistic will reduce by (*m* − 1)(*K*′ − 1) if these classes are aggregated, which will increase the power of the test if there are differences between the more abundant classes between environments. Aggregation of classes may be useful to prevent overfitting if the number of classes is large in comparison to the number of samples. The function to perform the Dirichlet likelihood ratio test given in the supplementary code has an option to include a *minimum proportion*. This is a value between 0 and 1 and is the minimum percentage threshold described above, divided by 100. For example, the 1% threshold used here would be described by a minimum proportion of 0.01. The default setting in the code is for the minimum proportion to be 0 and thus not be included.

The use of classes that are present across all samples out can be seen by considering the effect on (2) of having some *x*_*i*,*j*,*k*_ = 0. Specifically *L*(***α***|***x***) would be equal to 0 regardless of the choice of ***α***. As such we aggregate all classes that are not ever present. If this aggregated class still contains a zero sample then it is deleted and the data adjusted accordingly so that the compositions in each sample sum to 1. There is an option in the supplementary code to remove the need for ever present classes, which replaces all zeros with a small value (approximately 10^−16^) however, results from tests exercising this option should be quoted with extreme caution.

## Metagenomics ARG Discovery Methods

### Isolation of DNA from ovine interdigital swabs

Samples were collected from four sheep kept on the University of Nottingham, Sutton Bonington Campus farm during routine husbandry checks during the winter (November), spring (May) and summer (August) 2015/16. Sterile nylon flock swabs were used for the collection of samples from the interdigital space of sheep and stored in liquid amies solution at 5°C overnight. The swabs were processed following the methodology of (Frosth et al. 2015).

### DNA isolation and sequencing

DNA was isolated using the Qiagen Cador Pathogen Mini Kit, following the manufacturers guidelines and eluted in 60*μ*l of elution buffer. The DNA samples were quantified using the Qubit 3.0 and dsDNA high sensitivity dye. Quantified DNA was sent to Leeds Genomics (Leeds Univerity, U.K.) and prepared for sequencing using the NEB Library preparation kit and sequenced on the Illumina HiSeq 3000 at an approximate read depth of 50 million reads per sample.

### Analysis of sequence data

Raw reads were analysed for sequence adaptors using trimmomatic (Bolger et al. 2014) and clipped if necessary, the reads were then error corrected using the SGA k-mer based approach (Simpson and Durbin 2012). The corrected reads were firstly analysed using MetaPhlAn 2 (Truong et al. 2015) to identify bacteria present in the samples and then parsed against the MegaRes database (Lakin et al. 2017) using Diamond (Buchfink et al. 2015) to identify any antimicrobial resistance genes present. Finally BacMet (Pal et al. 2014) was used to identify any resistance genes associated to non-antibiotic elements.

## DATA ACCESS

Program code and data are available on FigShare at URL xxxxxxxx [to be completed upon acceptance for publication.]

## ACKNOWLEDGEMENTS

This work was supported by Engineering and Physical Sciences Research Council grants EP/P510993/1 and EP/M027333/1. We would also like to acknowledge the financial support by the National Natural Science Foundation of China (21210008) and K.C.Wong Education Foundation.

